# Direct visualization of bacterial transcripts in the infected lung illuminates spatiotemporal environmental adaptation of *Mycobacterium tuberculosis*

**DOI:** 10.1101/2025.08.25.672169

**Authors:** Anna-Lisa E. Lawrence, Shumin Tan

## Abstract

Spatiotemporal environmental variation results in marked heterogeneity in bacterial infection progression and disease outcome, with vital consequences for treatment success. For the globally important pathogen *Mycobacterium tuberculosis* (Mtb), while the pronounced intra-host spatial heterogeneity in lesion immune cell composition and phenotype has been well-described, the highly complex Mtb cell envelope has presented a particular challenge for the required equivalent insight into bacterial heterogeneity. Here, we develop hybridization chain reaction-fluorescence *in situ* hybridization (HCR-FISH)-based methodology for Mtb mRNA visualization in the context of intact lung and lesion architecture. In combination with a Mtb transcriptional/translational activity reporter, we reveal spatiotemporal differences in gene expression relating to Mtb lipid metabolism, response to key environmental signals, and the ESX-1 type VII secretion system. Our results establish a framework for *in* situ analysis of Mtb mRNA, opening the path to elucidating critical bacterial drivers that underlie the marked heterogeneity in Mtb-host interactions.

## INTRODUCTION

Bacterial adaptation to changes in the local environment is a critical driver of infection and disease outcome, with disruption of the bacterium’s ability to adapt and overcome host-mediated stressors resulting in colonization failure^1–7^. In addition to global changes in the host environment directed by factors such as the onset of adaptive immunity with time^8–11^, there has been a burgeoning appreciation for the spatial differences that also exist, facilitated by the development of techniques such as spatial transcriptomics, iterative protein or *in situ* hybridization-based microscopy, and imaging mass spectrometry-based methods^12–20^. This encompasses spatial differences ranging from host cell types and immune responses within lung or skin lesions during *Mycobacterium tuberculosis* (Mtb) or *Mycobacterium leprae* infection respectively^21–26^, to neutrophil infiltration and host cell metabolic changes in different regions of the corneal during *Pseudomonas aeruginosa* infection^27^, and to differences in availability of vital metals within the kidney during *Staphylococcus aureus* infection^20^. Such spatial heterogeneity in host cellular composition and response dictate a need for equivalent understanding of how the response and physiology of the infecting bacteria vary in tissue context, to elucidate specific bacterial niche adaptation responses and enable effective targeting of the bacteria for disease resolution.

To this end, fluorescent bacterial reporter strains have provided intriguing first insight into how bacterial gene expression changes at the single bacterium level during *in vivo* infection in spatial context. This spans studies on *Vibrio cholerae* toxin-related gene expression during intestinal infection^28^, the response of *Salmonella* and *Yersinia pseudotuberculosis* to nitric oxide stress in the spleen^29,30^, and intra-lesion differences in Mtb exposure to acidic pH and high chloride levels^31^. For Mtb, our studies with reporter strains have further revealed the correlation between a more acidic pH/higher chloride environment with decreased bacterial replication and activity, with a corresponding decrease in efficacy of drugs that target actively growing bacteria against Mtb residing in the lesion sublocation with more acidic pH/higher chloride^31^. Fluorescent reporters have thus been invaluable in uncovering key spatial changes in bacterial physiology during infection, but carry limitations such as restrictions in time points, in cases where the reporters are encoded on episomal plasmids, and in throughput. New methods that excitingly seek to enable more global probing of spatial changes in bacterial gene expression at the single bacterium level have more recently been reported, including the development of parallel and sequential FISH (par-seq FISH), which allowed for the detection of 105 different *P. aeruginosa* genes in planktonic and biofilm cultures^32^, and multiplexed error robust fluorescence *in situ* hybridization (MER-FISH), which coupled expansion microscopy with FISH-based labeling to detect individual bacterial mRNA in broth-grown *Escherichia coli*, and in *Bacteroides thetaiotaomicron* during colon colonization in spatial context^33^. Application of FISH-based techniques to the analysis of mRNA from bacteria with highly complex cell envelopes or thick peptidoglycan layers, which resist standard permeabilization methods and significantly impede probe access, however, presents an additional technical hurdle. A particular challenge is acid-fast Mtb, the causative agent of tuberculosis that remains the leading global cause of death from an infectious disease^34^, whose cell envelope comprises an outer capsule-like layer, mycolic acids, and arabinogalactan, peptidoglycan, and lipoarabinomannan layers beyond the plasma membrane^35^. At the same time, extensive studies revealing the spatial heterogeneity of host cellular architecture and response in hallmark Mtb lesions (granulomas) in the past few years^22–26^ emphasize the critical need to understand how Mtb response differs in spatial lesion context.

Here, we report the development of a method that enables direct, quantitative detection of individual Mtb operons via hybridization chain reaction-fluorescence *in situ* hybridization (HCR- FISH)^12,17^, at single bacterium resolution in the context of intact tissue and lesion architecture. We focus on three key facets of Mtb host infection (lipid utilization, nitric oxide (NO)/hypoxia exposure, and type VII secretion system function) to establish this method in our foundational study, utilizing the C3HeB/FeJ murine model of Mtb infection that recapitulates the hallmark necrotic lesion type found during human infection^36,37^. Our results reveal not just a temporal upregulation of Mtb core lipid-responsive genes as infection progresses, but spatial differences in expression of these genes between Mtb residing in the macrophage-dominant lesion cuff versus in the neutrophil-dominant core edge. Expression of the *hspX* operon that responds very robustly to initial exposure to NO and hypoxia showed striking differences in Mtb present in the lesion cuff versus in the core center, with the highest levels observed in the core edge. Finally, spatiotemporal differences in expression of the *espACD* ESX-1 type VII secretion system substrates were also observed, with highest expression in Mtb residing in the lesion cuff, and a trend towards reduced expression at the latest 16-week timepoint. Together with results from a chromosomally-encoded Mtb transcriptional/translational activity reporter, our work provides crucial insight into spatiotemporal changes in Mtb physiology and response during host infection, and raise intriguing concepts for future study. By establishing a framework and HCR-FISH methodology for analysis of Mtb mRNA, our study further opens the path to incorporating understanding of the bacterial aspect in elucidating factors that underlie the marked heterogeneity in Mtb-host interactions, enabling the holistic comprehension required to account for the critical impact of heterogeneity on infection outcome in designing effective therapeutic strategies.

## RESULTS

### Mtb activity within necrotic lesions decrease as infection progresses

Mtb growth state strongly impacts infection outcome and treatment efficacy^31,38–41^, yet spatiotemporal changes in Mtb replication in the context of necrotic lesion formation and maturation is not well understood. To address this question, we utilized a doxycycline-inducible monomeric Kusabira Orange (mKO) reporter Mtb strain that also carries a constitutively expressed mCherry (P_606_’::mKO-tetON, *smyc’*::mCherry), both encoded on the chromosome^31,42^. Active Mtb transcribe and translate mKO upon exposure of the mice to doxycycline inducer, enabling spatial analysis of which bacteria are active in different lesion sublocations, and we have shown that signal from this reporter tightly correlates with bacterial replication^31^. While we had previously found that penetration of doxycycline into the center of the lesion core (“core center”; Figure 1A) appears impeded as no mKO signal is observed, distinct differences between mKO signal induced upon doxycycline administration is observed between Mtb present in the lesion cuff versus in the edge of the lesion core (“core edge”) (Figure 1A)^31^. As a first examination of how Mtb activity changes spatiotemporally as infection progresses, we thus infected C3HeB/FeJ mice with Mtb(P_606_’::mKO-tetON, *smyc’*::mCherry) for 6, 12, or 16 weeks, followed by 1 week administration of doxycycline in the drinking water/food prior to sacrifice. Fixed lung samples were then analyzed for mKO signal in Mtb present in the lesion cuff versus core edge.

**Figure 1.**
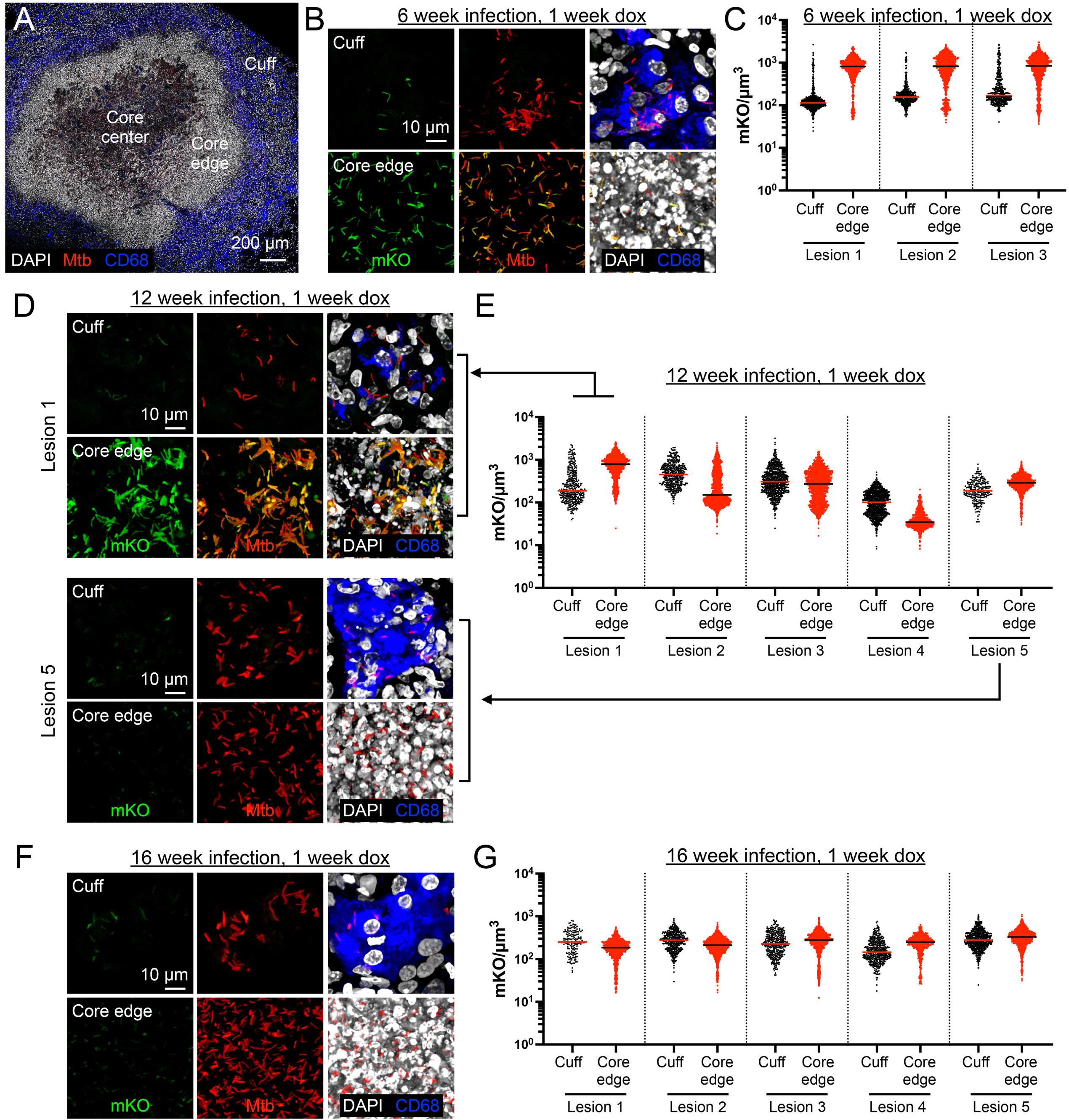
Mtb activity in the necrotic lesion core edge decreases as infection progresses. (A) Overview image of a necrotic lesion in C3HeB/FeJ mice at 7 wpi, with the lesion cuff, core edge, and core center marked. DAPI staining of nuclei is shown in grayscale, Mtb (*smyc’*::mCherry) in red, and CD68 staining of macrophages in blue. (B and C) Mtb activity is higher in the lesion core edge versus the cuff at 6 wpi. (B) shows representative 3D confocal images from a 6-week infection of C3HeB/FeJ mice with Mtb(P_606_’::mKO-tetON, *smyc’*::mCherry), followed by 1 week of exposure of the mice to doxycycline. All Mtb are marked in red (*smyc’*::mCherry), reporter signal in green (P_606_’::mKO-tetON), nuclei in grayscale (DAPI), and macrophages in blue (CD68). (C) shows quantification of mKO/µm^3^ signal for individual bacteria or a group of tightly clustered bacteria from 6-week infection + 1 week doxycycline-treated mice (3 lesions from 2 mice). Horizontal line marks the median. (D and E) Inter-lesion variability in Mtb transcriptional/translational activity profile at 12 wpi. (D) shows 3D confocal images from two examples of Mtb activity profiles from a 12-week infection of C3HeB/FeJ mice with Mtb(P_606_’::mKO-tetON, *smyc’*::mCherry), followed by 1 week of exposure of the mice to doxycycline. Staining and visualization are as in (B). (E) shows quantification of mKO/µm^3^ signal for individual bacteria or a group of tightly clustered bacteria from 12-week infection + 1 week doxycycline-treated mice (5 lesions from 4 mice). Horizontal line marks the median. (F and G) Mtb activity across lesion sublocations is decreased at 16 wpi. (F) shows representative 3D confocal images from a 16-week infection of C3HeB/FeJ mice with Mtb(P_606_’::mKO-tetON, *smyc’*::mCherry), followed by 1 week of exposure of the mice to doxycycline. Staining and visualization are as in (B). (G) shows quantification of mKO/µm^3^ signal for individual bacteria or a group of tightly clustered bacteria from 16-week infection + 1 week doxycycline-treated mice (5 lesions from 4 mice). Horizontal line marks the median.

Consistent with our previous results^31^, Mtb residing in the lesion core edge expressed high levels of mKO at 6 weeks post-infection (wpi) (Figures 1A-1C), a timepoint when such lesions first form^31,36^, indicating an environment favorable for bacterial replication. Mtb residing in the cuff, in contrast, expressed significantly lower levels of mKO at 6 wpi (Figures 1A-1C)^31^. There was notably increased heterogeneity between lesions at 12 wpi, with some lesions exhibiting similar patterns of Mtb mKO expression as those at 6 wpi (Figure 1D “lesion 1” and Figure 1E) (i.e. higher levels of mKO in Mtb residing in the core edge versus those in the cuff), while other lesions had lower levels of mKO signal in Mtb residing in both the cuff and core edge (Figure 1D “lesion 5” and Figure 1E). These results suggested a downward transition in Mtb activity levels, particularly of bacteria in the lesion core edge, as infection progressed. Indeed by 16 wpi, the patterns of Mtb activity levels in the lesion cuff versus core edge was consistent across lesions, with Mtb expressing low levels of mKO in both the lesion core edge and cuff (Figures 1F and 1G). The variability in Mtb mKO signal within each lesion sublocation was also markedly decreased at 16 wpi as compared to 6 wpi, with the coefficient of variation (CV) for Mtb in the lesion cuff decreasing to 54.95% ± 2.69% from 117.32% ± 9.21%, p<0.001, and to 45.27% ± 1.04% from 58.45% ± 3.16%, p<0.01, for Mtb in the lesion core edge (compare Figure 1G to Figure 1C).

These results indicate that Mtb activity decreases as lesions mature, with the lesion core edge becoming less favorable to bacterial replication as infection progresses.

### Adaptation of hybridization chain reaction fluorescence *in situ* hybridization (HCR-FISH) for detection of Mtb transcripts from *in vivo* samples

To understand how Mtb adapts to the changing environment as infection progresses and in spatial context, we pursued adaptation of hybridization chain reaction-fluorescence *in situ* hybridization (HCR-FISH) methodology to enable direct visualization of Mtb transcripts at the single bacterium level, in the context of intact tissue and lesion architecture. HCR-FISH is a quantitative approach where samples incubated with probes specific to an RNA of interest is proceeded by linear signal amplification via self-assembling fluorescently-labeled hairpins (Figure 2A)^12,17^. As a first test of HCR-FISH on Mtb infected lung samples, sectioned, paraffin-embedded lung tissue from C3HeB/FeJ mice infected for 13 weeks with Mtb constitutively expressing mCherry (*smyc’*::mCherry) were processed for analysis with probes against Mtb 16S ribosomal RNA (rRNA). Strong overlap between the 16S rRNA and mCherry signals was observed, supporting the specificity and utility of the 16S rRNA probes for marking all Mtb (Figure 2B). However, the complex cell envelope of Mtb^35^ presents a significant hurdle for detection of messenger RNA (mRNA) transcripts, which are much less abundant than rRNA^43–45^, with standard permeabilization techniques ineffective in enabling sufficient probe penetration for Mtb mRNA visualization. To this end, we developed a stepwise procedure to systematically disrupt the different layers of the Mtb cell envelope (Figure 2C, see Methods). First, deparaffinized lung sections were treated with amylase and pullulanase to digest Mtb capsule polysaccharides. This was followed by sequential treatment with sodium dodecyl sulfate (SDS) to disrupt envelope lipids, proteinase K and achromopeptidase to digest envelope proteins, and hydrochloric acid to disrupt the mycolic acid layer. Subsequently, treatment with recombinantly expressed and purified trehalose dimycolate (TDM) hydrolase from *Mycobacterium smegmatis*^46^ and mycobacteriophage protein lysin B^47^ target TDM and mycolylarabinogalactan, with a final lysozyme treatment for disruption of the peptidoglycan layer (Figure 2C). Importantly, these treatments allowed for the detection of gene operons in Mtb (described below), while not affecting the ability to stain for host nuclei with DAPI, nor the robust detection of Mtb 16S rRNA.

**Figure 2.**
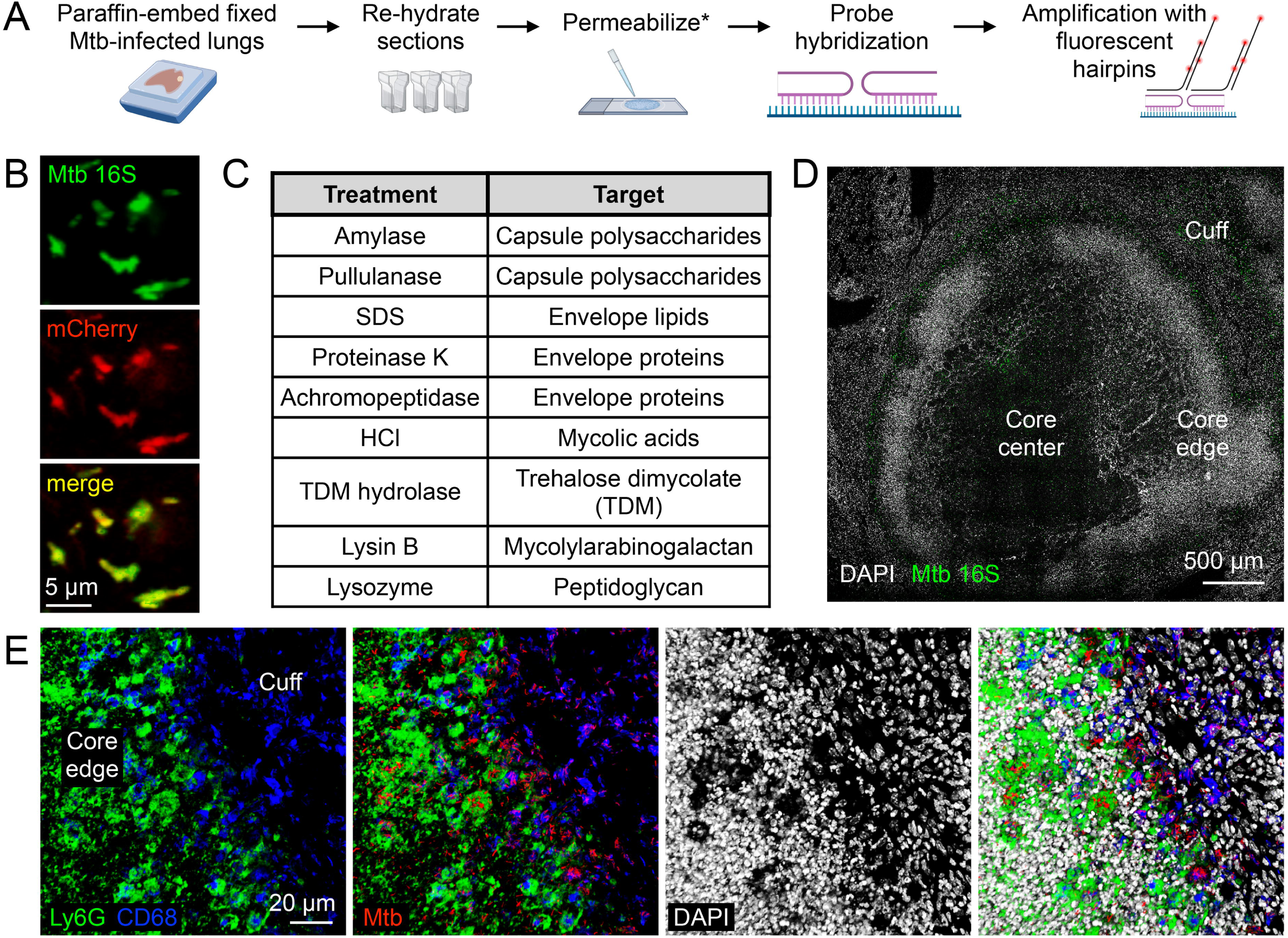
Adaptation of HCR-FISH for detection of Mtb transcripts in the context of intact lung tissue. (A) Schematic depicting workflow of HCR-FISH for Mtb transcripts. Generated using BioRender (https://BioRender.com/qfa1hd4). (B) HCR-FISH probes against Mtb 16S rRNA effectively label all Mtb. 13 wpi mCherry Mtb-infected C3HeB/FeJ lung sample was probed with 16S rRNA (green) with HCR-FISH and imaged via confocal microscopy. (C) List of permeabilization treatments used to enable detection of Mtb mRNA, with the targets for each respective treatment noted. (D) Lesion overview at 6 wpi highlighting the 3 lesion sublocations analyzed for expression of different Mtb operons using HCR-FISH. Mtb 16S rRNA is shown in green and host nuclei stained with DAPI is shown in grayscale. (E) The lesion cuff is macrophage-dominant and the lesion core edge is neutrophil-dominant. 3D confocal images of the interface region of the lesion cuff and core edge from a 6 wpi mCherry Mtb-infected C3HeB/FeJ lung sample is shown. Ly6G (neutrophil) staining is shown in green, CD68 (macrophage) in blue, and DAPI staining of nuclei in grayscale. All Mtb are shown in red (*smyc’*::mCherry).

We focused our studies here on 3 timepoints from C3HeB/FeJ mice infections to capture spatiotemporal changes in Mtb transcriptional responses over the course of infection: (i) 2 weeks, which is prior to lesion formation, (ii) 6 weeks, when canonical necrotic lesions are first routinely detected^31,36^, and (iii) 16 weeks, which our studies with the inducible mKO reporter show significant downregulation of Mtb activity compared to earlier timepoints. For spatial analysis, responses were categorized for Mtb residing in the lesion cuff (predominantly intracellularly within macrophages), in the lesion core edge (a mixture of Mtb present within neutrophils and among necrotic cell debris), and in the lesion core center (extracellular Mtb) (Figures 2D and 2E)^31,36^. Notably, in a subset of lesions at 16 wpi, 16S rRNA signal was significantly weaker than the signal observed at earlier time points (Figure S1), suggesting that these bacteria may be in the process of being cleared by the host. For robustness of comparison, analysis of mRNA expression in the 16 wpi samples was restricted to those lesions that had similar levels of Mtb 16S rRNA expression as the earlier time points.

### Spatiotemporal changes in expression of Mtb lipid utilization genes

Lipids, specifically cholesterol and fatty acids, are a critical nutrient source for Mtb and mutant Mtb strains that cannot utilize lipids are attenuated *in vivo*^6,48,49^. To determine where and when Mtb expresses genes involved in lipid response, indicating exposure to and usage of lipids as a carbon source, we designed probes for *rv3160c*-*rv3162c*, part of the core lipid response of Mtb^50,51^. At 2 wpi, Mtb resides primarily intracellularly within macrophages, and at this time point, low or non-detectable levels of *rv3160c-rv3162c* was observed in most bacteria (97.6% ± 1.2%), with very few, if any, bacteria expressing intermediate or high levels of this operon (Figures 3A, 3D, and S2A). Lipid-rich foamy macrophages are not observed at this early timepoint^6^, and this result directly supports that there is little exposure of Mtb to lipids as a primary carbon source at this early stage of infection.

**Figure 3.**
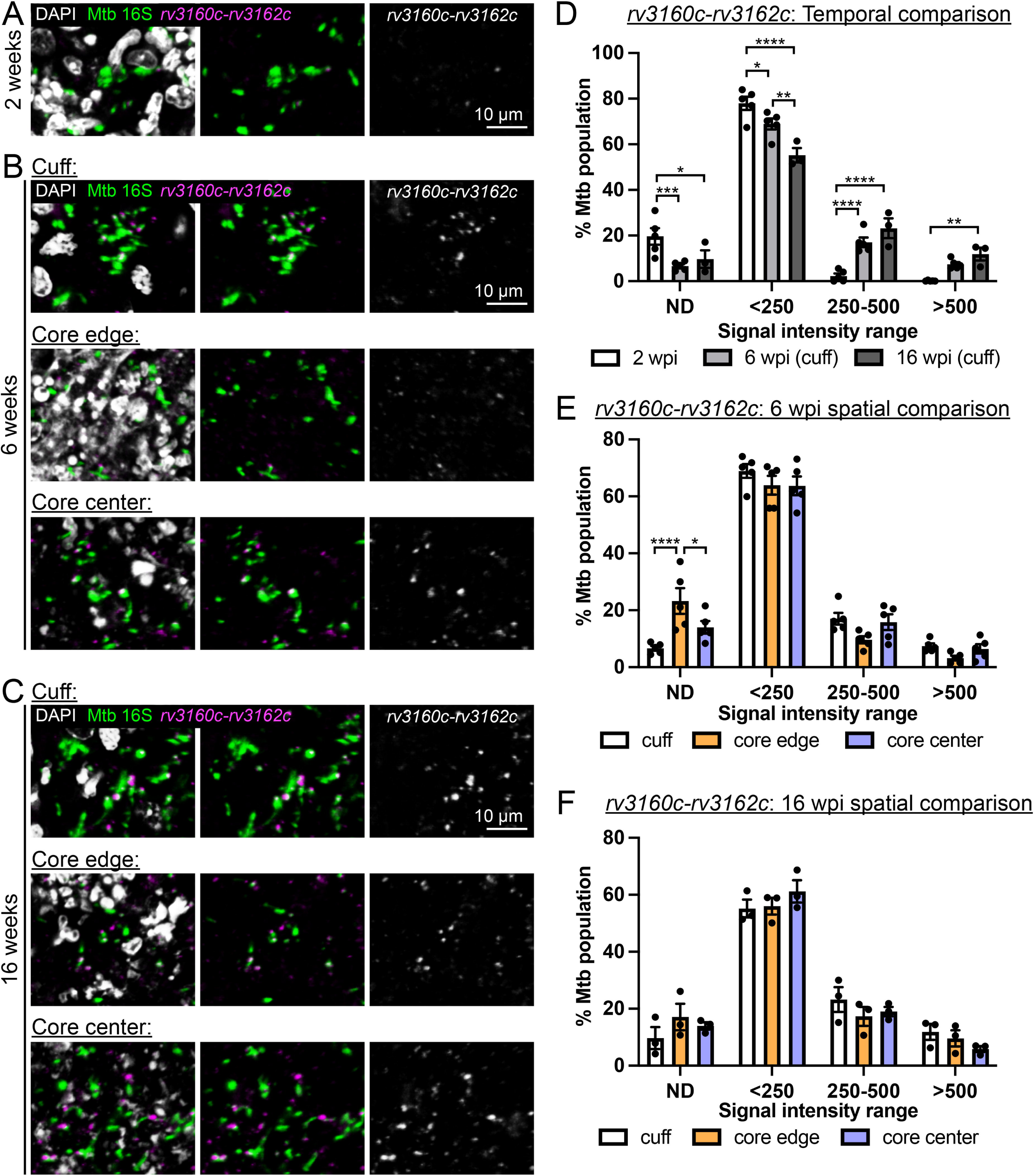
Spatiotemporal changes in expression of Mtb lipid utilization genes. (A-C) Representative 3D confocal images of Mtb at 2 wpi (A), 6 wpi (B), and 16 wpi (C) in C3HeB/FeJ mice, probed for the core lipid response genes *rv3160c-rv3162c* (magenta). Mtb 16S rRNA labels all bacteria (green) and DAPI staining of host nuclei is shown in grayscale. Merged image is shown on the left, merged image without DAPI signal in the middle panel, and *rv3160c- rv3162c* signal is shown alone in the right panel (in grayscale for clarity). (D) *rv3161c-rv3162c* expression increases with time post-infection. The percentage of Mtb expressing different levels of *rv3160c-rv3162c* at 2 wpi and from the lesion cuff region at 6 and 16 wpi is shown, separated into 4 bins – non-detectable (ND), low (<250 signal/µm^3^), intermediate (250-500 signal/µm^3^), or high (>500 signal/µm^3^). (E and F) Mtb in the lesion core edge express lower levels of *rv3160c- rv3162c* at 6, but not 16, wpi. The percentage of Mtb in each lesion sublocation expressing different levels of *rv3160c-rv3162c* at 6 (E) and 16 (F) wpi is shown, separated into 4 bins – non-detectable (ND), low (<250 signal/µm^3^), intermediate (250-500 signal/µm^3^), or high (>500 signal/µm^3^). Data are from 5 mice for 2 wpi, from 5 lesions from 5 mice for 6 wpi, and from 3 lesions from 2 mice for 16 wpi. p-values were obtained with a 2-way ANOVA with Tukey’s multiple comparisons test in (D) – (F). Only significant comparisons are indicated. * p<0.05, ** p<0.01, *** p<0.001, **** p<0.0001.

At 6 wpi, *rv3160c-rv3162c* expression markedly increased in Mtb present in the lesion cuff (bacteria primarily intracellular in macrophages) (Figure 3B, top row), with a significant increase in the percentage of Mtb expressing intermediate (250-500 signal intensity/µm^3^) and high (>500 signal intensity/µm^3^) levels of *rv3160c-rv3162c* (24.0% ± 2.6% combined, versus 2.4% ± 1.2% at 2 wpi, p<0.001) (Figures 3B, 3D, 3E, and S2B). A marked increase was similarly observed for Mtb present in the lesion core center (22.3% ± 4.2%) (Figures 3B and 3E). The necrotic core has previously been reported to be lipid-rich, due to high levels of necrotic host cells making up the caseum, including lipid-rich foamy macrophages^52^. Interestingly, a greater percentage of Mtb present in the lesion core edge still had non-detectable levels of *rv3160c- rv3162c* transcript (23.2% ± 4.5% versus 6.6% ± 0.8% for the cuff, p<0.0001; and versus 14.0% ± 2.2% for the core center, p<0.05), with a correspondingly decreased percentage showing intermediate and high levels (Figures 3B, 3E, and S2B). Finally, a high percentage of Mtb in all sublocations of necrotic lesions expressed *rv3160c-rv3162c* at 16 wpi, with a trend of increased expression compared to the 6 wpi samples (Figures 3C, 3D, 3F, and S2C).

Together, these results reveal an overall transition of Mtb from a non-lipid-rich to lipid-rich local environment as necrotic lesions form. They further demonstrate how the dynamic nature of differences in Mtb localization alters its exposure to lipids, with spatial differences in the lesion core edge versus center during the early stages of lesion development that dissipate at later stages.

### Nitric oxide/hypoxia early responsive genes are upregulated by Mtb residing in the lesion cuff and core edge

As well as adaptation to changing nutrient sources, Mtb response to environmental cues vitally modulate bacterial growth. Nitric oxide (NO) and hypoxia represent two critical environmental cues able to drive Mtb into an adaptive non-replicating state, with upregulation of a set of genes known as the dormancy regulon^53–55^. The two-component system DosRS(T) regulates the dormancy regulon, with the *hspX* operon most often used as a marker of the DosR- dependent hypoxia/NO-mediated transcriptional response^53–58^. To gain insight into where and when Mtb encounters these critical stressors during infection, HCR-FISH probes were thus designed against the *hspX* operon genes. Expression of the *hspX* operon was detected even at 2 wpi, with only 9.5% ± 2.8% of Mtb with non-detectable probe signal (Figures 4A, 4D, and S3A). This was initially surprising, as we had previously observed much greater expression of *hspX* at 4 versus 2 wpi in C57BL/6 mice using a *hspX’*::GFP reporter Mtb strain^59^. To follow up this finding, we performed HCR-FISH on the same infected tissue to test for expression of *Nos2*, which encodes inducible nitric oxide synthase (iNOS), the enzyme responsible for producing NO during Mtb infection^60^. Consistent with Mtb *hspX* operon expression at 2 wpi, *Nos2* was detected in the lungs of infected C3HeB/FeJ mice at this timepoint, with robust expression comparable to that observed in 6 wpi Mtb-infected lung samples (Figure S4).

**Figure 4.**
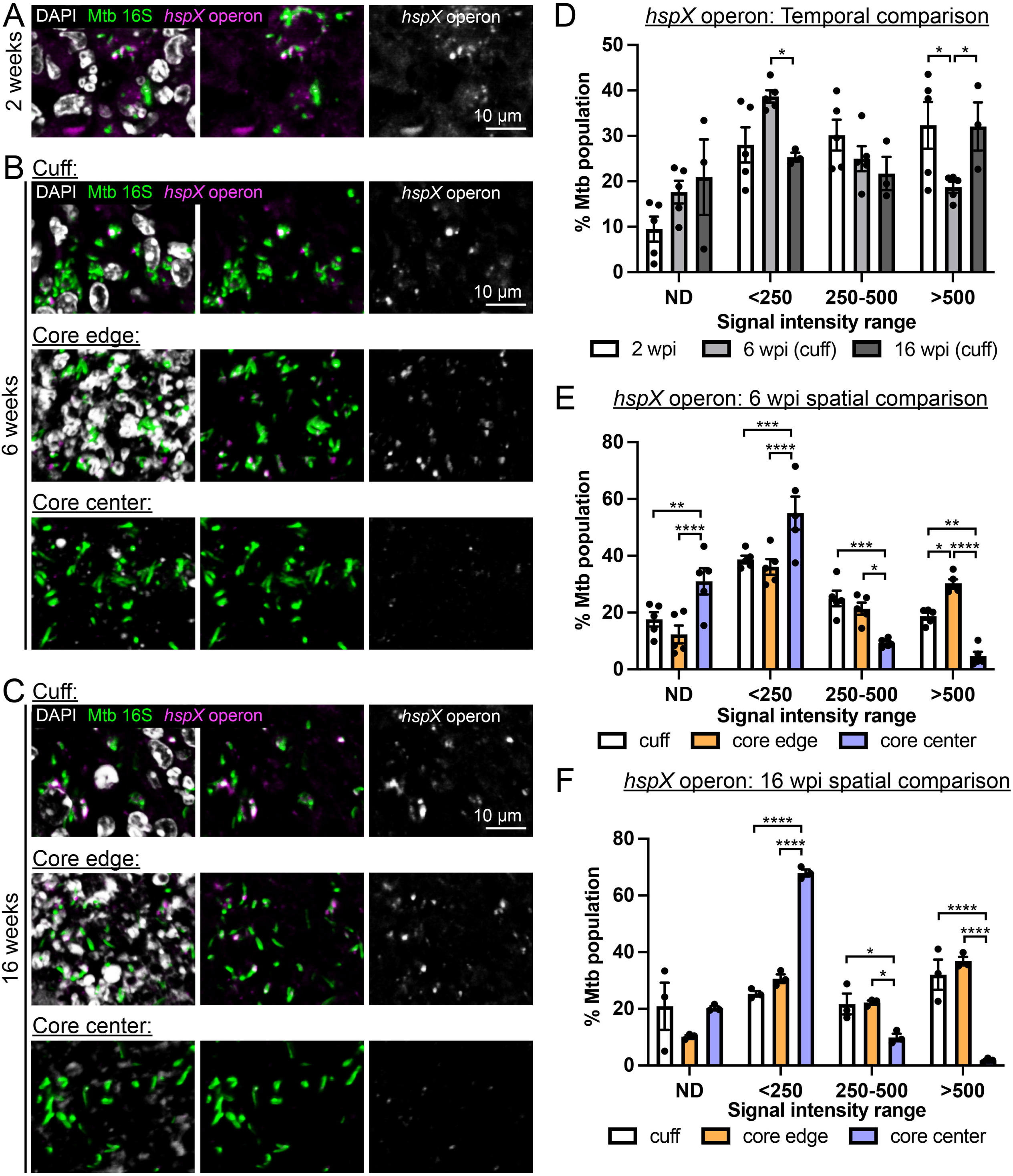
**Nitric oxide/hypoxia early responsive genes are upregulated by Mtb residing in the lesion cuff and core edge**. (A-C) Representative 3D confocal images of Mtb at 2 wpi (A), 6 wpi (B), and 16 wpi (C) in C3HeB/FeJ mice, probed for the *hspX* operon (magenta). Mtb 16S rRNA labels all bacteria (green) and DAPI staining of host nuclei is shown in grayscale. Merged image is shown on the left, merged image without DAPI signal in the middle panel, and *hspX* operon signal is shown alone in the right panel (in grayscale for clarity). (D) The *hspX* operon is expressed throughout infection in Mtb residing in macrophages. The percentage of Mtb expressing different levels of the *hspX* operon at 2 wpi and from the lesion cuff region at 6 and 16 wpi is shown, separated into 4 bins – non-detectable (ND), low (<250 signal/µm^3^), intermediate (250-500 signal/µm^3^), or high (>500 signal/µm^3^). (E and F) Mtb in the lesion core edge express higher levels of the *hspX* operon at 6, but not 16, wpi, with expression very low in the lesion core center. The percentage of Mtb in each lesion sublocation expressing different levels of the *hspX* operon at 6 (E) and 16 (F) wpi is shown, separated into 4 bins – non-detectable (ND), low (<250 signal/µm^3^), intermediate (250-500 signal/µm^3^), or high (>500 signal/µm^3^). Data are from 5 mice for 2 wpi, from 5 lesions from 5 mice for 6 wpi, and from 3 lesions from 2 mice for 16 wpi. p-values were obtained with a 2-way ANOVA with Tukey’s multiple comparisons test in (D) – (F). Only significant comparisons are indicated. * p<0.05, ** p<0.01, *** p<0.001, **** p<0.0001.

At 6 and 16 wpi, robust *hspX* operon expression was also observed in a significant percentage of Mtb residing in the lesion cuff, with 34.3% ± 3.4% and 51.9% ± 8.8% combined, respectively, expressing intermediate (250-500 signal intensity/µm^3^) or high (>500 signal intensity/µm^3^) levels (Figures 4B-4D and S3B-S3C). Intriguingly, the percentage of Mtb expressing the highest levels of the *hspX* operon was even greater in the Mtb population residing in the lesion core edge versus the lesion cuff at 6 wpi (30.3% ± 1.5% versus 18.7% ± 1.2%, p<0.05) (Figure 4E). As neutrophils produce NO and further consume oxygen in generation of the oxidative burst^61,62^, it is possible that this finding reflects the presence of both NO and hypoxia in this lesion sublocation. This difference between the lesion cuff and core edge is however lost at 16 wpi, with the percentage of Mtb expressing the highest levels of the *hspX* operon in the lesion cuff increasing to now match that observed in the lesion core edge (32.1% ± 5.3% versus 36.9% ± 1.5%) (Fig. 4F).

Strikingly, we found that the *hspX* operon was expressed at significantly lower levels in the necrotic core center, with only 4.7% ± 1.5% and 1.9% ± 0.4% of bacteria in this sublocation at 6 and 16 wpi, respectively, exhibiting the highest signal levels (Figures 4B-4C, 4E-4F, and S3B-S3C). Importantly, although the *hspX* operon responds robustly to hypoxia, this initial strong response is not sustained, with the *hspX* operon not part of the characterized “enduring hypoxia response”^63^. The high percentage (68.0% ± 1.2%) of Mtb residing in the necrotic lesion core with a low level (<250 signal intensity) of *hspX* expression at 16 wpi might thus reflect an extended exposure of that subpopulation of Mtb to hypoxia.

Together, these results show that Mtb is exposed to NO-related stress as early as 2 wpi in this infection model. The spatiotemporal differences in expression of the *hspX* operon in necrotic lesions further reveals how Mtb residence in different immune cells or extracellularly alters its exposure to the vital environmental cues of NO and hypoxia.

### ESX-1 Type VII secretion system effector genes are expressed most strongly by Mtb present in the lesion cuff and decreases with time

Finally, in addition to undergoing shifts in metabolism and responding to environmental cues during infection, Mtb directly manipulates interactions with host immune cells via systems such as the ESX-1 type VII secretion system^64–66^. The ESX-1 type VII secretion system has been shown to be critical in Mtb infection^64–68^, yet the timing and spatial regulation of these genes remains largely unknown. To examine expression of this critical virulence factor spatiotemporally, we thus designed probes against the *espACD* operon, which encode 3 important substrates of the ESX-1 secretion system^69–72^. The majority of bacteria exhibited *espACD* expression even at 2 wpi, with just 9.7% ± 3.8% of Mtb with non-detectable levels (Figures 5A, 5D, and S5A), supporting the importance of the ESX-1 type VII secretion system in host colonization by Mtb.

**Figure 5.**
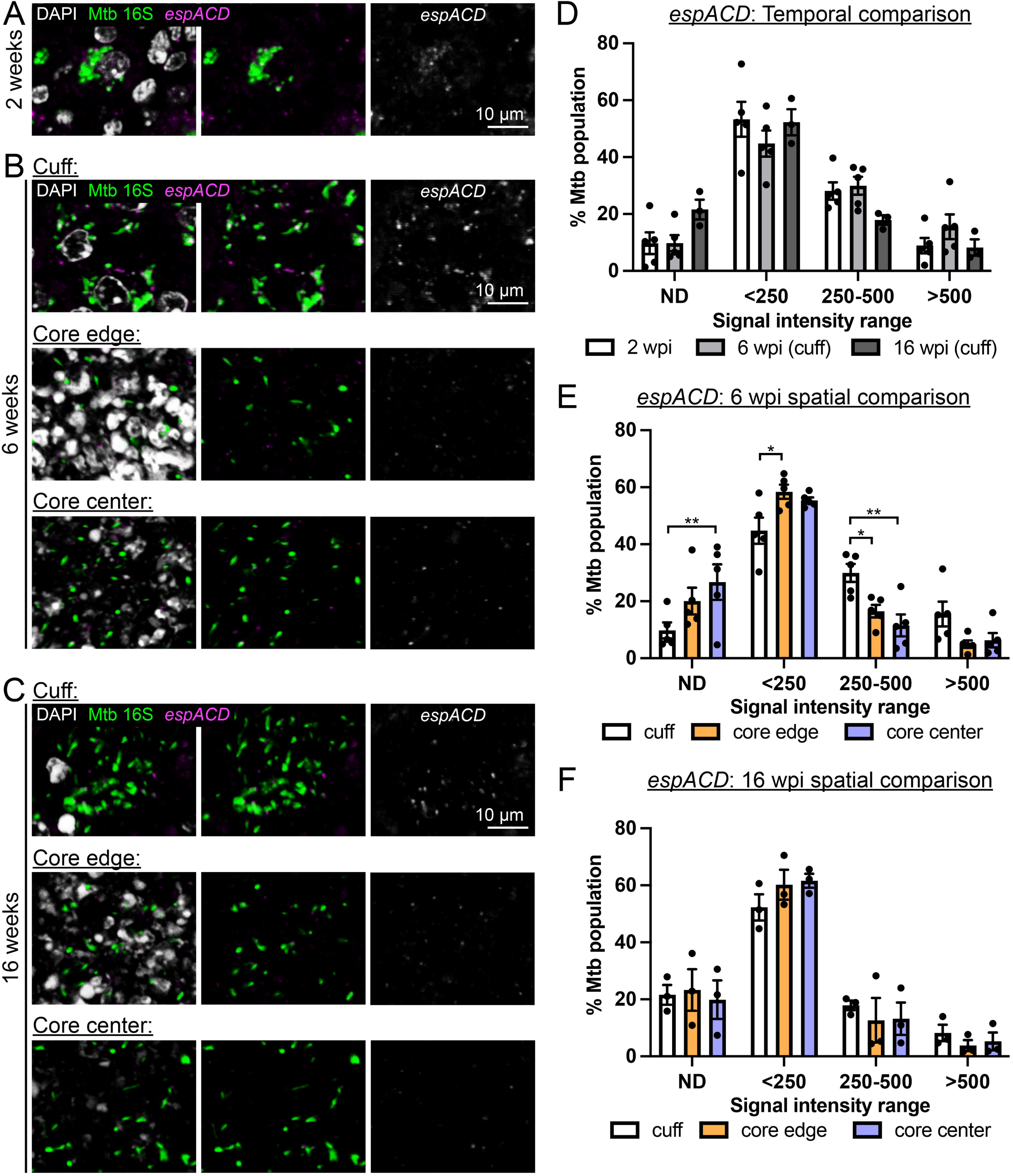
ESX-1 Type VII secretion system effector genes are expressed most strongly by Mtb present in the lesion cuff and decreases with time. (A-C) Representative 3D confocal images of Mtb at 2 wpi (A), 6 wpi (B), and 16 wpi (C) in C3HeB/FeJ mice, probed for the *espACD* operon (magenta). Mtb 16S rRNA labels all bacteria (green) and DAPI staining of host nuclei is shown in grayscale. Merged image is shown on the left, merged image without DAPI signal in the middle panel, and *espACD* operon signal is shown alone in the right panel (in grayscale for clarity). (D) The *espACD* operon is expressed beginning early in infection and trends down at 16 wpi. The percentage of Mtb expressing different levels of the *espACD* operon at 2 wpi and from the lesion cuff region at 6 and 16 wpi is shown, separated into 4 bins – non- detectable (ND), low (<250 signal/µm^3^), intermediate (250-500 signal/µm^3^), or high (>500 signal/µm^3^). (E and F) Mtb in the lesion cuff express higher levels of the *espACD* operon at 6, but not 16, wpi. The percentage of Mtb in each lesion sublocation expressing different levels of the *espACD* operon at 6 (E) and 16 (F) wpi is shown, separated into 4 bins – non-detectable (ND), low (<250 signal/µm^3^), intermediate (250-500 signal/µm^3^), or high (>500 signal/µm^3^). Data are from 5 mice for 2 wpi, from 5 lesions from 5 mice for 6 wpi, and from 3 lesions from 2 mice for 16 wpi. p-values were obtained with a 2-way ANOVA with Tukey’s multiple comparisons test in (D) – (F). Only significant comparisons are indicated. * p<0.05, ** p<0.01.

Expression of *espACD* in Mtb residing in the lesion cuff was similarly robust at 6 wpi, with just 9.8% ± 2.8% of Mtb with non-detectable levels (Figures 5B, 5D-5E, and S5B). In contrast, at 6 wpi, *espACD* expression levels were significantly lower in Mtb residing in the lesion core edge, and particularly in the lesion core center, with a smaller percentage expressing intermediate or high levels of this operon versus Mtb residing in the lesion cuff (21.5% ± 3.3% and 17.9% ± 6.3% combined for the core edge and core center, respectively, versus 45.5% ± 6.7% for the cuff, p<0.05 in each case) (Figures 5B-5C, 5E, and S5B). Most bacteria in the lesion core edge and core center continued to express low or non-detectable levels of *espACD* at 16 wpi (Figures 5C, 5F, and S5C). Interestingly, at 16 wpi, there was a downward trend of *espACD* expression in Mtb resident in the lesion cuff compared to 6 wpi, with 21.6% ± 3.5% of the population in this lesion sublocation now with non-detectable levels (versus 9.8% ± 2.8% at 6 wpi, p<0.05) (Figures 5E-5F and S5B-S5C).

Together, these findings provide key insight into the spatiotemporal expression of the ESX-1 type VII secretion system, indicating its importance particularly earlier during infection for Mtb residing in macrophages.

## DISCUSSION

While the impact of heterogeneity during Mtb infection on disease outcome and treatment efficacy is now well-appreciated, with multiple studies employing single cell technologies focused on understanding host cell differences in spatiotemporal context^22–26^, studies analyzing the bacterial aspect have posed a continued significant technical challenge. The use of reporter strains and techniques such as laser capture microdissection combined with bulk RNA sequencing have been invaluable in providing first insight into Mtb physiology during infection^31,57,58,73,74^. However, reporter strains that are encoded on episomal plasmids are limited to shorter-term infections and in their throughput, while laser capture microdissection-RNA sequencing approaches do not provide single cell resolution. More recently, FISH-based methods have been applied to Mtb but have been restricted to either highly expressed RNA (rRNA) or used many probes (120) together against several unrelated genes for Mtb mRNA detection^75–78^. Our establishment here of HCR-FISH methodology for direct detection of single operons in individual Mtb in the context of intact lung tissue opens the path to spatiotemporally studying Mtb responses at the single bacterium level.

The spatial differences observed in our analysis of three key aspects of Mtb biology (lipid, NO/hypoxia, type VII secretion) in three distinct lesion sublocations (cuff, core edge, core center), enabled by the single bacterium resolution afforded by HCR-FISH, raise several intriguing questions for follow-up study. First, it highlights distinct differences in Mtb resident in macrophages (lesion cuff) versus in neutrophils (lesion core edge). This is particularly the case when lesions have first formed (6 wpi), with lower expression levels of the lipid core response genes *rv3160c-rv3162c* and the ESX-1 type VII secretion system substrates *espACD*, and higher levels of the *hspX* operon, in Mtb residing in the lesion core edge versus cuff. While neutrophils can accumulate lipids^79–81^, how these levels compare to those in foamy macrophages and whether access of Mtb to lipids present in neutrophils versus in foamy macrophages differs is unknown. Mtb exhibits slowed growth when utilizing lipids as a carbon source^82,83^, and the lower levels of *rv3160c-rv3162c* expression of Mtb in the lesion core edge versus the cuff at 6 wpi is thus in accord with the higher Mtb activity in this sublocation revealed by the inducible mKO reporter experiment. While higher *hspX* operon expression appears counterintuitive to higher Mtb activity, it is important to note both that a similar phenomenon of active Mtb in neutrophils despite high *hspX’*::GFP reporter and inducible nitric oxide expression has previously been observed in an early infection model of C57BL/6J mice^84^, and that the mKO range of Mtb in the lesion core edge at 6 wpi is very large, with clear bifurcation into active and inactive Mtb subpopulations in some cases^31^. It will be interesting in future studies to multiplex readouts, to directly examine how *hspX* operon expression relates to Mtb activity and growth.

Considering the spatial differences in *espACD* expression, while ESX-1-dependent neutrophil necrosis *in vitro* has been reported^85^, the role of ESX-1 in Mtb-neutrophil interactions *in vivo* is largely unknown. The release of neutrophil extracellular traps (NETs) in a manner that does not result in neutrophil death and supports Mtb growth was recently discovered, and interestingly, strong staining of citrullinated histone 3 (NET marker) was observed in the region between the macrophage-rich lesion cuff and the caseous center in Mtb-infected cynomolgus macaque lung samples^86^. Notably, recent studies have begun to delineate the existence of different neutrophil subsets during Mtb infection that have differing impact on Mtb^79,87–89^. Mtb- neutrophil studies to date have largely focused on *in vitro* and early infection timepoints in animal models; as both Mtb growth status and utilization of lipids as a carbon source impact critically on treatment efficacy^31,38–41,90,91^, our findings here further draw attention to the need to understand how Mtb interaction with its host cell differs depending on host cell type/subtype, and between and within lesion sublocations, for effective therapeutic regimen design. Understanding how newly discovered neutrophil subsets may map onto the lesion context and the differences in Mtb physiology observed here will thus be important to examine in future studies.

Second, the striking low levels of *hspX* operon expression in Mtb present in the lesion core center compared to bacilli in the lesion cuff or core edge highlight the very different local environment experienced by extracellular Mtb in the lesion core center. The core center of mature lesions has always been assumed to be hypoxic, even if direct determination has been technically precluded by the need for live host cells for positive pimonidazole labeling^92–94^. While the *hspX* operon is widely used in the field as a marker of Mtb response to NO and hypoxia given its extremely strong upregulation upon initial exposure (on the order of hours) to these signals^53–55^, it is crucial to note that this upregulation decreases with time. Indeed, a different set of genes has been described to mark an extended response to hypoxia^63^, and further studies examining members of this “enduring hypoxia response” gene set will be needed to robustly define how oxygen availability within the lesion core center changes as infection progresses.

Temporally for bacteria residing in macrophages, our results provide direct bacterial response data supporting the switch to lipid metabolism as infection progresses, in line with foamy macrophage formation as necrotic lesions form. At the same time, the increase in *hspX* operon and decrease in *espACD* signals for Mtb residing in the lesion cuff from 6 to 16 wpi raise questions as to whether there is a transition of the macrophage subtype in which the bacteria reside as infection progresses. Interstitial macrophages are known to be more restrictive for Mtb growth versus alveolar macrophages^84^, but how they may differentially contribute to lesion composition in spatiotemporal context, as well as the provenance of the abundant foamy macrophages found in the necrotic lesion cuff, is not well-understood. The significant decrease in variation in Mtb activity status in the lesion cuff with time (decrease in coefficient of variation of the inducible mKO signal) further points to changes in Mtb-host cell interplay as infection progresses and lesions mature. Our finding of a downward trend in expression of the type VII secretion system substrates *espACD* with time reinforces the need to understand spatiotemporal changes in bacterial responses in treatment design, particularly as therapy must successfully target established, not just beginning, infections, for effectiveness against a chronic disease like tuberculosis.

The HCR-FISH methodology for Mtb mRNA detection presented here enables a straightforward and accessible approach for testing the expression of specific Mtb operons of interest in spatial tissue/lesion context. We propose that future studies building on the foundational framework established here will continue to provide crucial insight into what drives Mtb population heterogeneity during infection and facets vital to bacterial survival and adaptation to specific local niches. For example, multiplexing different Mtb operons that reflect the bacterial response to different signals, with host aspects, or with use of the FISH-based “RS ratio” (ratio of short-lived spacer rRNA region to stable mature rRNA, which serves as a proxy for rRNA synthesis activity that tightly correlates with bacterial replication^75,77,95^), will allow direct *in situ* analysis of the relationships between Mtb environmental response, growth, and host cell/location phenotype. Utilization of this approach in combination with drug treatment or specific perturbation of Mtb regulatory pathways further holds immense promise in revealing how drug efficacy is affected spatiotemporally, and how targeting of key nodes in Mtb signal integration can be leveraged for altering infection heterogeneity to aid treatment success. Finally, the HCR-FISH approach established here can also be used to directly probe tissue samples from patients, potentially opening a path to the study not just of the most common pulmonary manifestation of Mtb infection, but also disease outcomes with poor animal models. We additionally anticipate that this approach can be broadly adapted to other difficult to permeabilize bacteria for which FISH-based detection of mRNA have also posed technically challenging, such as *Streptococcus pneumoniae* and *Clostridiodes difficile*, to excitingly enable interrogation of the impact of spatiotemporal heterogeneity across multiple other important infectious diseases.

## ACKNOWLEDGEMENTS

We thank Graham Hatfull and Anil Ojha for the kind gift of plasmids for recombinant expression and purification of lysin B and TDM hydrolase, respectively. We thank members of the Tan lab for helpful discussion. This work was supported by grants to ST from the Hypothesis Fund and from the National Institutes of Health (R01 AI143768). The funders had no role in study design, data collection and analysis, decision to publish, or preparation of the manuscript.

## AUTHOR CONTRIBUTIONS

Conceptualization: AEL, ST

Methodology: AEL, ST

Formal analysis: AEL, ST

Investigation: AEL, ST

Writing – original draft: AEL, ST

Writing – review and editing: AEL, ST

Supervision: ST

Funding acquisition: ST

## STAR METHODS RESOURCE AVAILABILITY

### Lead contact

Further information and requests for reagents should be directed to and will be fulfilled by the lead contact, Shumin Tan.

### Materials availability

The reporter *M. tuberculosis* strain used in this study will be shared as requested with investigators with the necessary biosafety level 3 facilities to receive and work with this material.

### Data and code availability

All data reported in this paper will be shared by the lead contact upon request. This paper does not report original code. Any additional information required to reanalyze the data reported in this paper is available from the lead contact upon request.

## EXPERIMENTAL MODEL AND SUBJECT DETAILS

### Mtb strains and culture

Mtb Erdman wild type was used in this study. Edman(P_606_’::mKO-tetON, *smyc’*::mCherry) has been previously described^31^. Preparation of mouse infection stocks was as previously described^57^.

### Mice

Animal protocols were reviewed and approved by the Institutional Animal Care and Use Committee at Tufts University (#B2024-90), in accordance with the Association for Assessment and Accreditation of Laboratory Animal Care, the US Department of Agriculture, and the US Public Health Service guidelines. These protocols followed standards set by the National Institutes of Health “Guide for the Care and Use of Laboratory Animals”. 6 week old female C3HeB/FeJ wild type mice (Jackson Laboratory, Bar Harbor, ME) were used for mouse infection studies.

## METHOD DETAILS

### Mouse Mtb infections

C3HeB/FeJ wild type mice (Jackson Laboratory, Bar Harbor, ME) were intranasally infected with 10^3^ colony forming units of Mtb in 35 µl of phosphate-buffered saline (PBS) containing 0.05% Tween-80, under light anesthesia with 2% isoflurane^31,57,58^. Mice were sacrificed at 2, 6, 7, 13, 16, or 17 wpi and lungs fixed in 4% paraformaldehyde (PFA) in PBS overnight at room temperature, then stored in phosphate buffered saline (PBS) supplemented with 100 U

SUPERase·In RNase inhibitor (ThermoFisher Scientific) prior to further processing. For the P_606_’::mKO-tetON, *smyc’*::mCherry infections, one week prior to sacrifice, either drinking water containing 1 mg/ml doxycycline with 5% sucrose or food containing 200 mg/kg doxycycline (Bioserv) were supplied to the mice^31,96^.

### Recombinant protein expression and purification

Plasmids encoding TDM hydrolase or lysin B^46,47^ were expressed in *E. coli* BL21(DE3) in the presence of 100 µg/ml ampicillin. 1 L cultures in LB broth were grown to OD_600_ ∼0.6 prior to induction with 1 mM IPTG for 3 hours at 30°C (TDM hydrolase) or 4 hours at 37°C (lysin B). Cells were pelleted and resuspended in low imidazole buffer (500 mM NaCl, 50 mM Tris pH 7.5, 15 mM imidazole, 10% glycerol) before being flash-frozen in liquid nitrogen and stored at - 80°C prior to purification. Thawed cells were lysed by sonication, in the presence of protease inhibitors (Pierce mini protease inhibitor tablets, ThermoFisher Scientific). The soluble fraction was collected and incubated overnight with 1 ml nickel-NTA agarose beads (Machery-Nagel) at 4°C, with agitation. To elute the protein, beads were washed 3x with low imidazole buffer prior to elution in high imidazole buffer (500 mM NaCl, 50 mM Tris pH 7.5, 200 mM imidazole, 10% glycerol). Samples were dialyzed overnight at 4°C to remove imidazole using Slide-A-Lyzer G3 dialysis cassettes (ThermoFisher Scientific) in 50 mM Tris pH 7.5, 100 mM NaCl, 25 mM MgCl_2_ (TDM hydrolase), or in 50 mM Tris pH 8, 50 mM NaCl (lysin B)^46,47^. Protein was collected from the dialysis cassettes the next day, aliquoted, flash-frozen in liquid nitrogen, and stored at -80°C until use.

### HCR-FISH on Mtb transcripts

For HCR-FISH analyses, lung samples were processed for paraffin embedding and sectioning (5 µm thick sections) (Tufts Comparative Pathology Services) and stored at 4°C in air-tight containers with desiccant until use. Paraffin was removed by washing with xylene substitute (Fisher Scientific) for 3 x 5 minutes, followed by rehydration using decreasing concentrations of ethanol from 100% to 50% for 3 minutes each, with a final wash step in UltraPure water (ThermoFisher Scientific) for 3 minutes. Samples were then permeabilized to detect Mtb transcripts, via sequential treatment in a humidified chamber with: (i) 1 µg/ml amylase (MilliporeSigma) and 1:1000x dilution of pullulanase (MilliporeSigma) in 20 mM sodium acetate, pH 5 buffer for 1 hour at 37°C; (ii) 1% SDS in PBS for 15 minutes at room temperature; (iii) 10 µg/ml proteinase K (Fisher Scientific) in Tris HCl pH 7 for 10 minutes at 37°C; (iv) 30 U achromopeptidase (MilliporeSigma) in Tris HCl pH 8.5 for 40 minutes at 37°C; (v) 0.2 M HCl for 2.5 minutes at 37°C; (vi) TDM hydrolase and lysin B at 1 mg/ml and 1.5 mg/ml respectively in 50 mM Tris pH 8, 100 mM NaCl and 25 mM MgCl_2_ for 2 hours at 37°C; and (vii) 10 mg/ml lysozyme (Roche) in TE buffer pH 7 for 1 hour at 37°C. All treatments were carried out in 100 µl volumes, and 200 U SUPERase·In RNase inhibitor (ThermoFisher Scientific) was added to inhibit RNase activity during permeabilization. Following permeabilization, antigen retrieval was performed in Tris-EDTA buffer at 95°C for 15 minutes, after which slides were cooled to 45°C in 20 minutes by adding 40 ml UltraPure water every 5 minutes. Slides were further incubated in 50 ml UltraPure water at room temperature for 10 minutes prior to 2 x 2 minute washes with PBS containing 0.1% Tween-20 (PBST).

For probe hybridization, the prepared tissue slices were first incubated with 200 µl probe hybridization buffer (Molecular Instruments) with 0.1 mg/ml salmon sperm DNA (ThermoFisher Scientific) for 10 minutes prior to addition of probes at 0.02 µM concentration overnight in probe hybridization buffer at 37°C in a humidified chamber. Probes were designed against Mtb 16S rRNA (labels all bacteria), *rv3160c*-*rv3162c* (lipid-responsive), the *hspX* operon ((*rv2028c*- *rv2031c*; hypoxia/nitric oxide-responsive), and the *espACD* (*rv3614c-rv3616c*) operon (type VII secretion system substrates). The following day, unbound probes were removed by washing with decreasing concentrations of probe wash buffer (Molecular Instruments) (100% to 0%) diluted in 5x saline-sodium citrate buffer containing 0.1% tween-20 (SSCT) at 37°C for 15 minutes each, followed by one wash at room temperature with 5x SSCT for 5 minutes. Tissue sections were then treated with 200 µl amplification buffer containing 0.1 mg/ml salmon sperm DNA for 30 minutes at room temperature. At the same time, 4 µl of 3 µM hairpin stock per slide were heated at 95°C for 90 seconds in a thermocycler and then cooled down to RT, protected from light, for 30 minutes. To detect 16S rRNA in tissue, Alexa Fluor 488 and 647 amplifiers were used for 16S rRNA and mRNA target detection, respectively. Amplifier fluorophores in amplification buffer were added directly to the tissue samples prior to overnight incubation at room temperature. The next day, slides were first washed with 5x SSCT for 5 minutes, then again with 5x SSCT for 2 x

15 minutes, with a final 5x SSCT wash for 5 minutes. Samples were stained with DAPI (ThermoFisher Scientific) diluted 1:500 in blocking buffer (3% BSA + 0.1% triton X-100 in PBS) for 10 minutes at room temperature. Slides were washed 3x with 5x SSCT prior to mounting with Prolong Glass (ThermoFisher Scientific). Coverslips were allowed to cure overnight at room temperature prior to imaging.

### HCR-FISH on host transcripts

To detect expression of *Nos2* in Mtb-infected lungs, paraffin-embedded sections were processed and stained as described above, but excluding the additional permeabilization steps required for

Mtb permeabilization. Briefly, after tissue rehydration, lung sections underwent antigen retrieval at 95°C for 15 minutes prior to allowing the samples to cool. Slides were treated with 2 x 2 minute PBST washes followed by probe hybridization. Signal amplification was performed as described for Mtb transcripts.

### Confocal microscopy and image quantification

For the P_606_’::mKO-tetON, *smyc’*::mCherry reporter infections, lung lobes were embedded in 4% agarose in PBS and 250 µm thick sections obtained using a Leica VT1000S vibratome^31,97^. Lung sections were blocked with 3% BSA + 0.1% Triton X-100 in PBS (blocking buffer) for 1 hour at room temperature (all steps protected from light) before overnight incubation with rat anti-CD68 (Bio-Rad) primary antibody at 1:100 dilution in blocking buffer at room temperature. The following day, lung samples were washed 3 x 5 min with blocking buffer prior to staining with Alexa Fluor 647 goat anti-rat secondary antibody (ThermoScientific; 1:100 dilution) and DAPI (ThermoScientific; 1:500 dilution) for 2 hours at room temperature in blocking buffer. Tissue slices were mounted using VECTASHIELD mounting medium (Vector labs). Samples were imaged on a Leica SP8 spectral confocal microscope using a 40x oil immersion objective. z-stacks were 10 µm thick with 0.5 µm z-steps. Images were reconstructed into 3D using Volocity software (Quorum Technologies, Inc). HCR-FISH samples were imaged using a Leica SP8 spectral confocal microscope with z-stacks that were 3 µm thick with 0.35 µm z-steps. For all microscopy analyses, regions were selected based on the presence of bacteria without regard to reporter or mRNA signal, and lesions across multiple mice and multiple experiments sampled.

Quantification of mKO reporter and mRNA signal was performed as previously described using Volocity software^31,57,58,97^. Briefly, Mtb volume was determined via either mCherry signal (for the mKO reporter infection) or 16S rRNA signal (for HCR-FISH studies) and the total signal (either mKO or mRNA probes of interest) within each bacterium measured. Settings for mKO and each mRNA target were maintained across samples to allow for direct comparison across lesions and timepoints. For HCR-FISH analysis, normalization for differences in background signal was also performed, with the background fluorescence intensity/µm^3^ of 8 random locations not containing bacteria measured and subtracted from the HCR-FISH signal for each image.

## QUANTIFICATION AND STATISTICAL ANALYSIS

Details of data quantification and statistical analysis utilized are described in the figure legends. Statistical tests were computed using GraphPad Prism software. A 2-way ANOVA with Tukey’s multiple comparisons test was utilized in Figures 3D-3F, 4D-4F, and 5D-5F. p<0.05 was considered significant in all cases.

## KEY RESOURCES TABLE

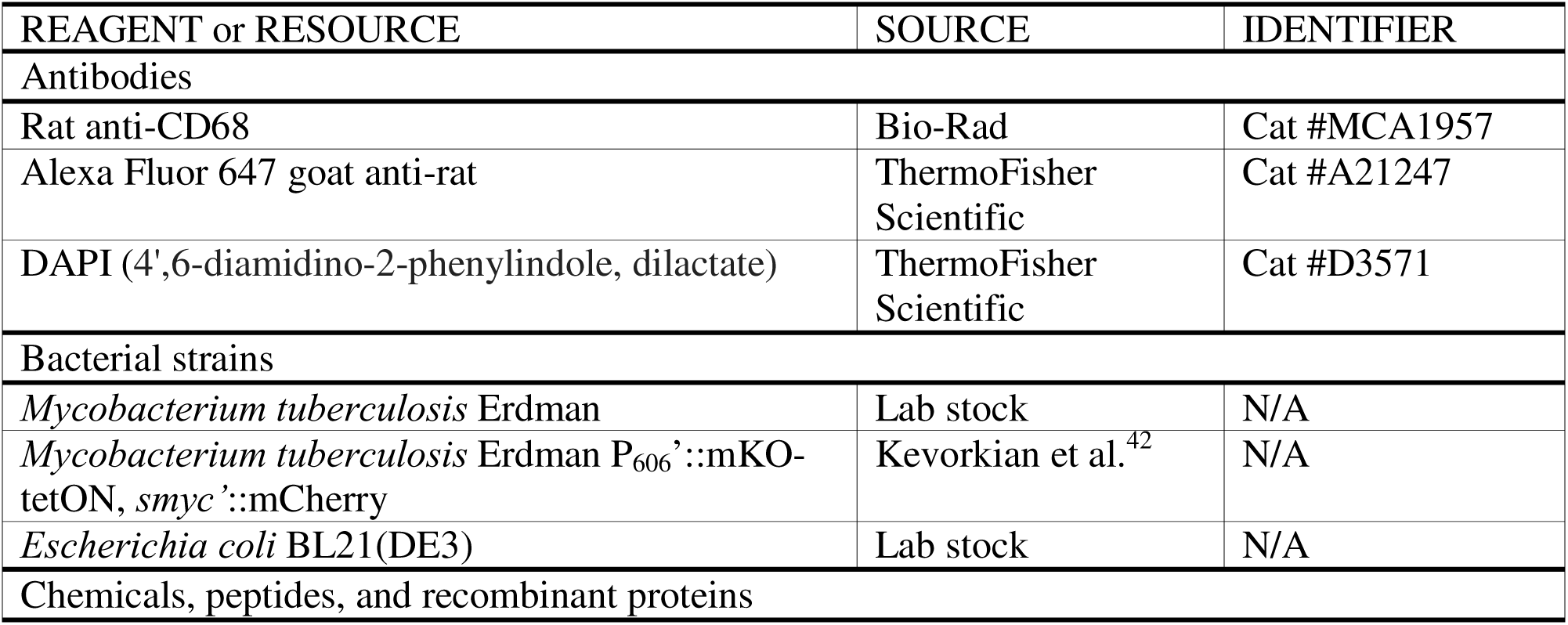
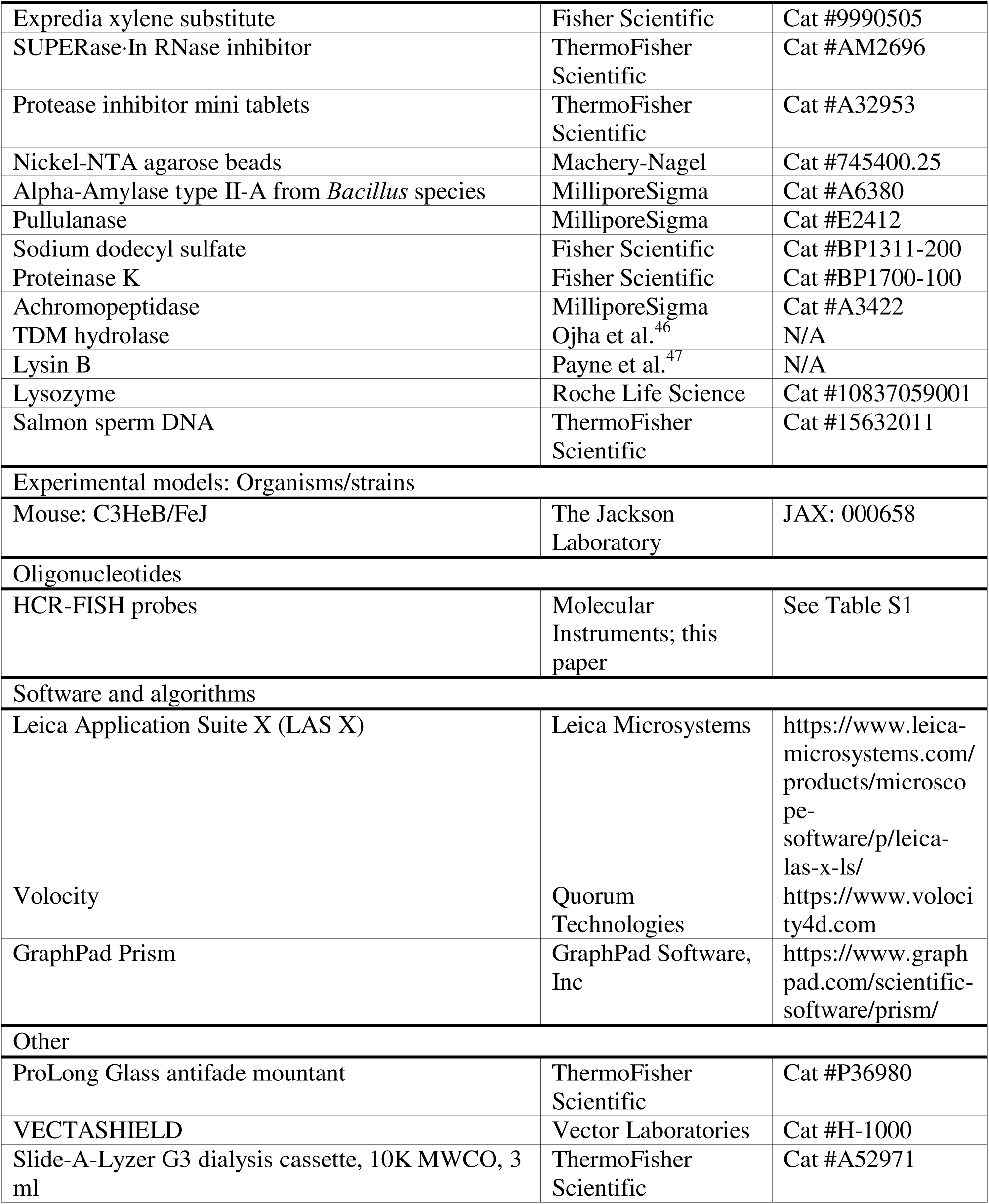

## SUPPLEMENTAL FIGURE LEGENDS

**Figure S1. Mtb in a subset of 16 wpi lesions exhibit lower 16S rRNA levels.** C3HeB/FeJ mice were infected with Mtb for 16 weeks and lung tissues processed for HCR-FISH. Mtb 16S rRNA signal/µm^3^ for individual bacteria or a group of tightly clustered bacteria, from 5 different lesions across 3 mice, were analyzed in each of the different lesion sublocations.

**Figure S2. Quantification of *rv3160c-rv3162c* expression in individual bacteria.** *rv3160c- rv3162c* signal/µm^3^ for individual bacteria or a group of tightly clustered bacteria at 2 (A), 6 (B), or 16 (C) wpi in C3HeB/FeJ mice. Horizontal lines mark the median for each sample, with the percentage of bacteria with non-detectable (“ND”) levels of expression indicated at the bottom of each sample. For 6 and 16 wpi timepoints when necrotic lesions have formed, Mtb residing in the three lesion sublocations (cuff, core edge, or core center) were separately analyzed. Data are from 5 mice for 2 wpi, from 5 lesions from 5 mice for 6 wpi, and from 3 lesions from 2 mice for 16 wpi.

**Figure S3. Quantification of *hspX* operon expression in individual bacteria.** *hspX* operon signal/µm^3^ for individual bacteria or a group of tightly clustered bacteria at 2 (A), 6 (B), or 16 (C) wpi in C3HeB/FeJ mice. Horizontal lines mark the median for each sample, with the percentage of bacteria with non-detectable (“ND”) levels of expression indicated at the bottom of each sample. For 6 and 16 wpi timepoints when necrotic lesions have formed, Mtb residing in the three lesion sublocations (cuff, core edge, or core center) were separately analyzed. Data are from 5 mice for 2 wpi, from 5 lesions from 5 mice for 6 wpi, and from 3 lesions from 2 mice for 16 wpi.

**Figure S4. N*os2* expression is observed in host cells even at 2 wpi.** 3D confocal images of 2 (top row) or 6 (bottom row) wpi lung Mtb-infected C3HeB/FeJ lung samples analyzed for *Nos2* expression (yellow) and Mtb 16S rRNA (green) via HCR-FISH. Nuclei (DAPI) are shown in grayscale.

**Figure S5. Quantification of *espACD* operon expression in individual bacteria.** *espACD* operon signal/µm^3^ for individual bacteria or a group of tightly clustered bacteria at 2 (A), 6 (B), or 16 (C) wpi in C3HeB/FeJ mice. Horizontal lines mark the median for each sample, with the percentage of bacteria with non-detectable (“ND”) levels of expression indicated at the bottom of each sample. For 6 and 16 wpi timepoints when necrotic lesions have formed, Mtb residing in the three lesion sublocations (cuff, core edge, or core center) were separately analyzed. Data are from 5 mice for 2 wpi, from 5 lesions from 5 mice for 6 wpi, and from 3 lesions from 2 mice for 16 wpi.

