## Supplementary figures and images for "Direct visualization of bacterial transcripts in the infected lung illuminates spatiotemporal environmental adaptation of *Mycobacterium tuberculosis*"

### Figure S1

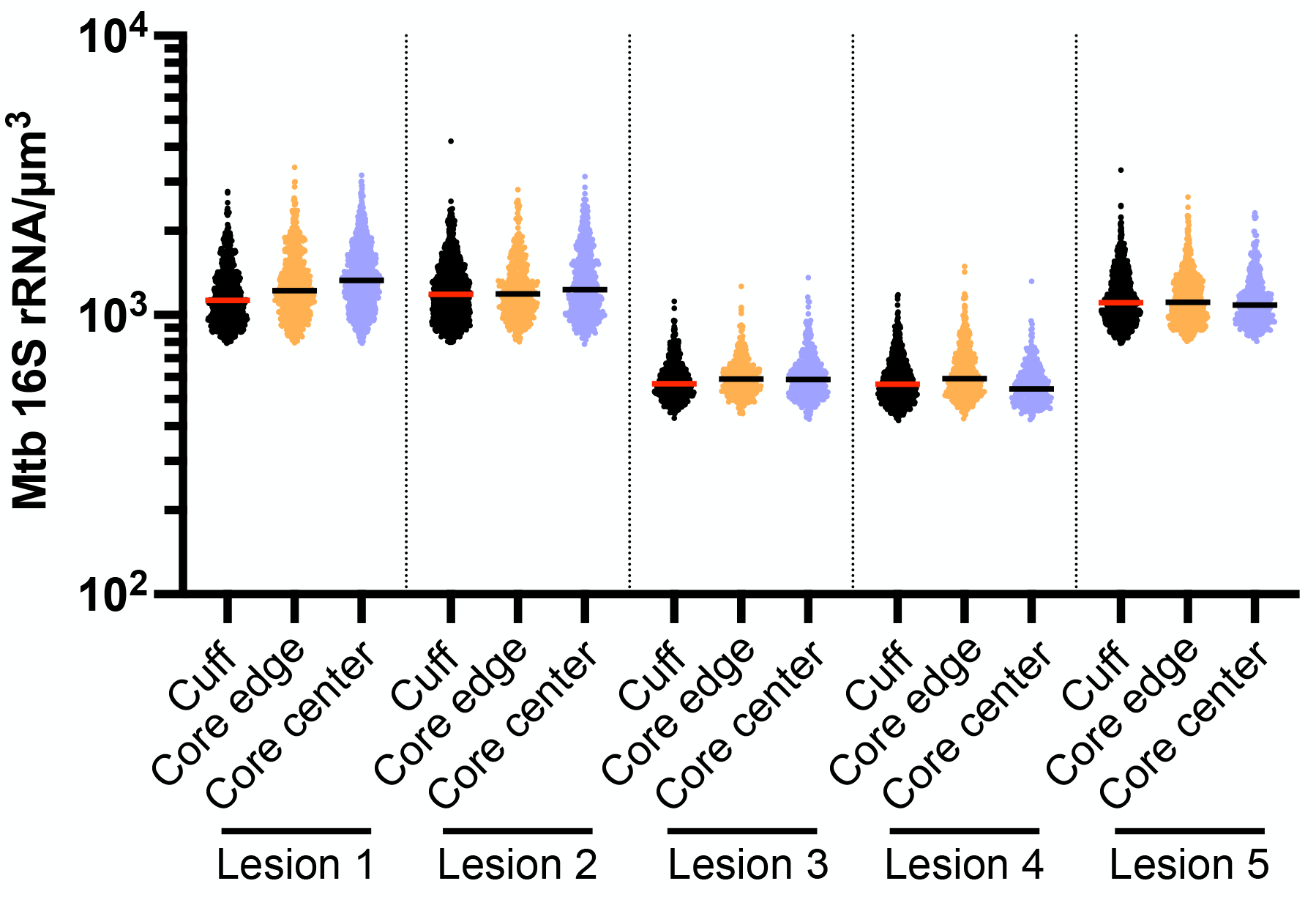

### Figure S2

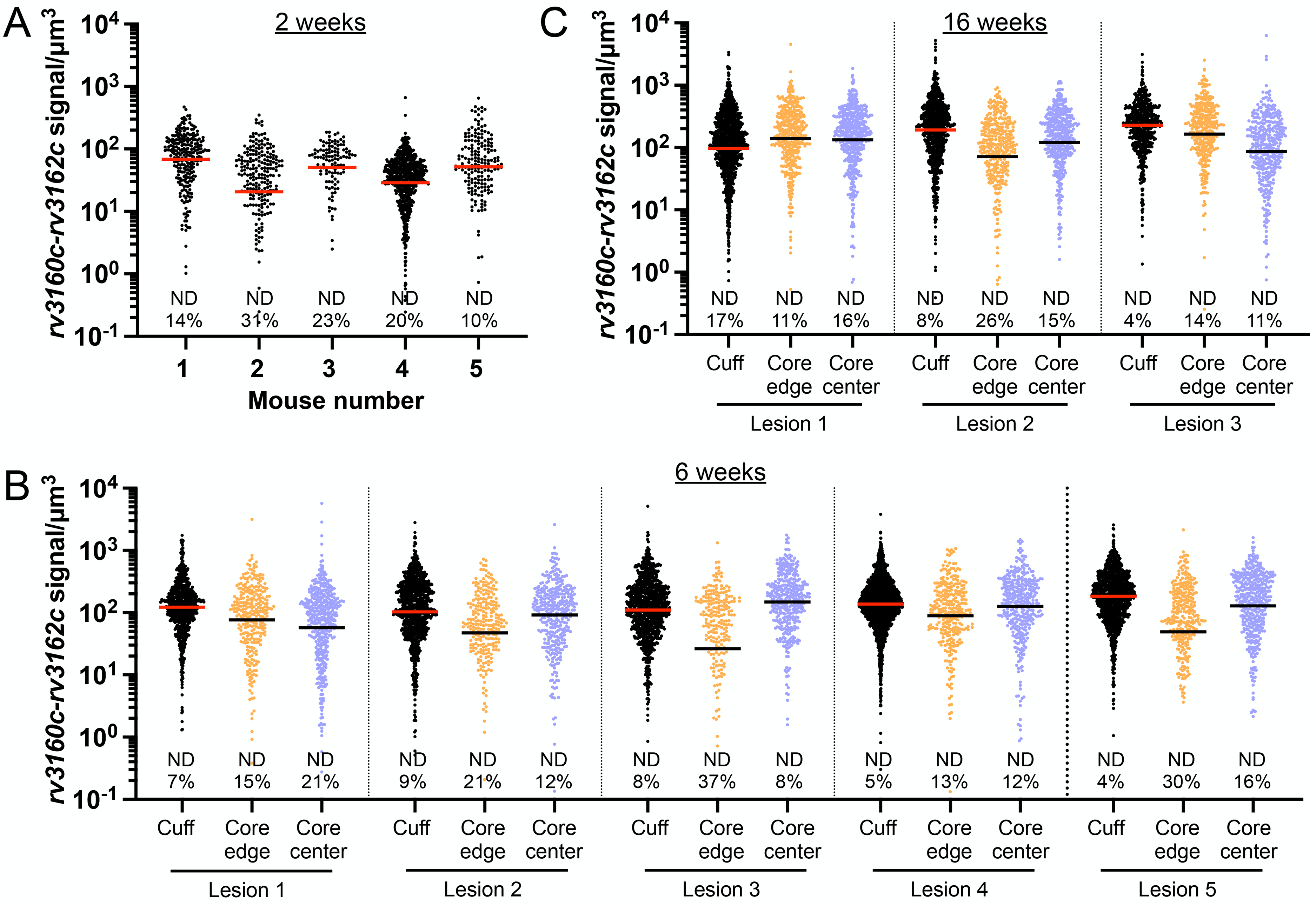

### Figure S3

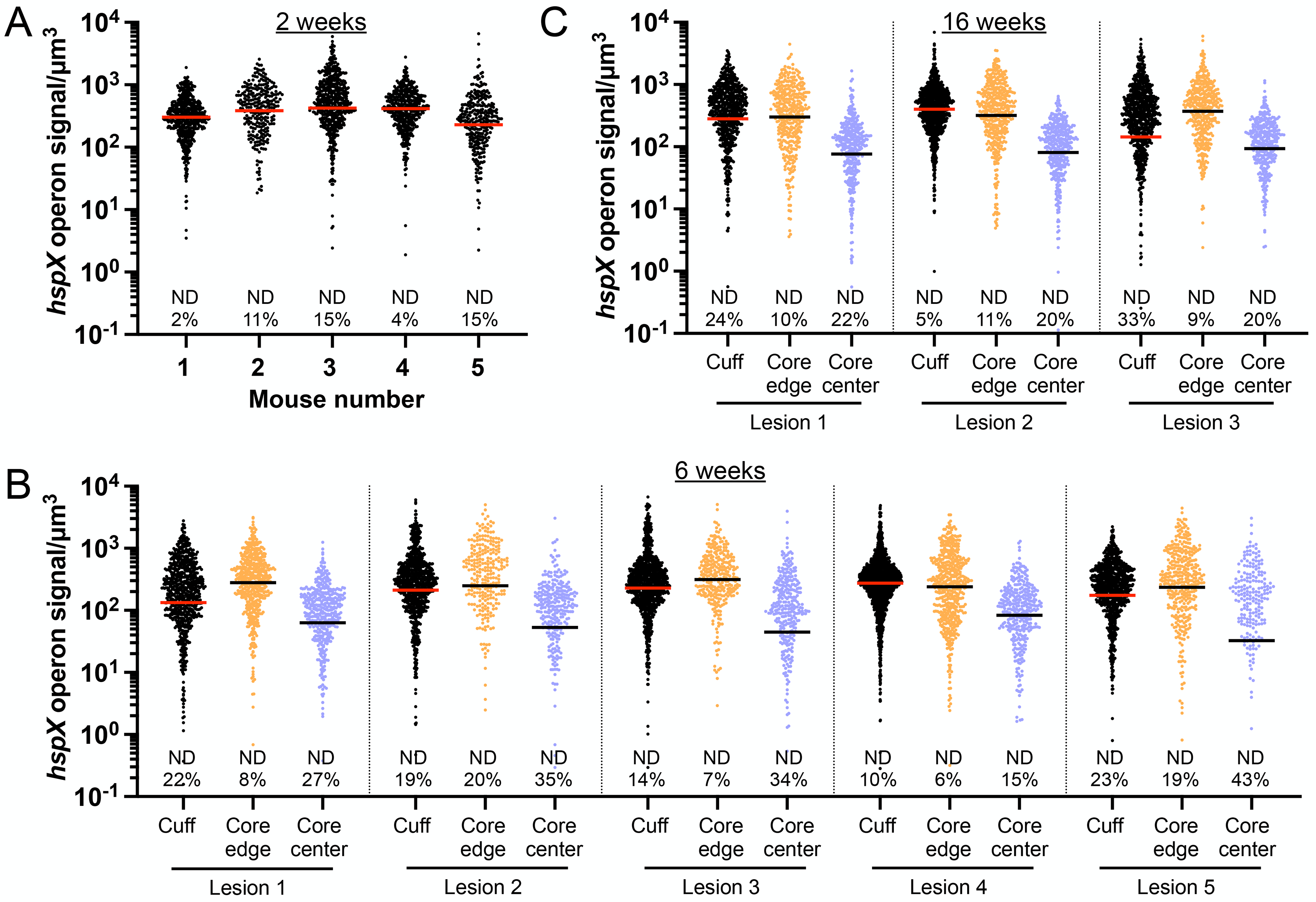

### Figure S4

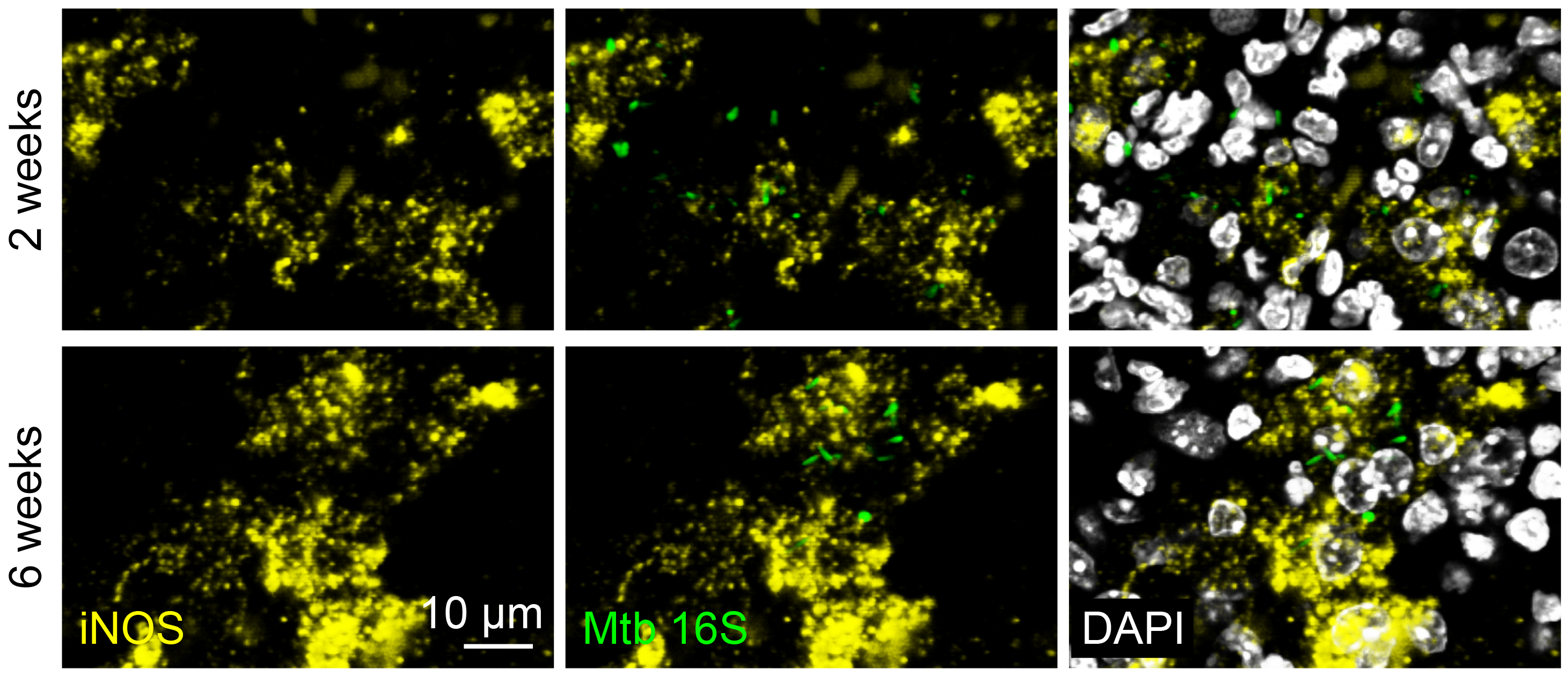

### Figure S5

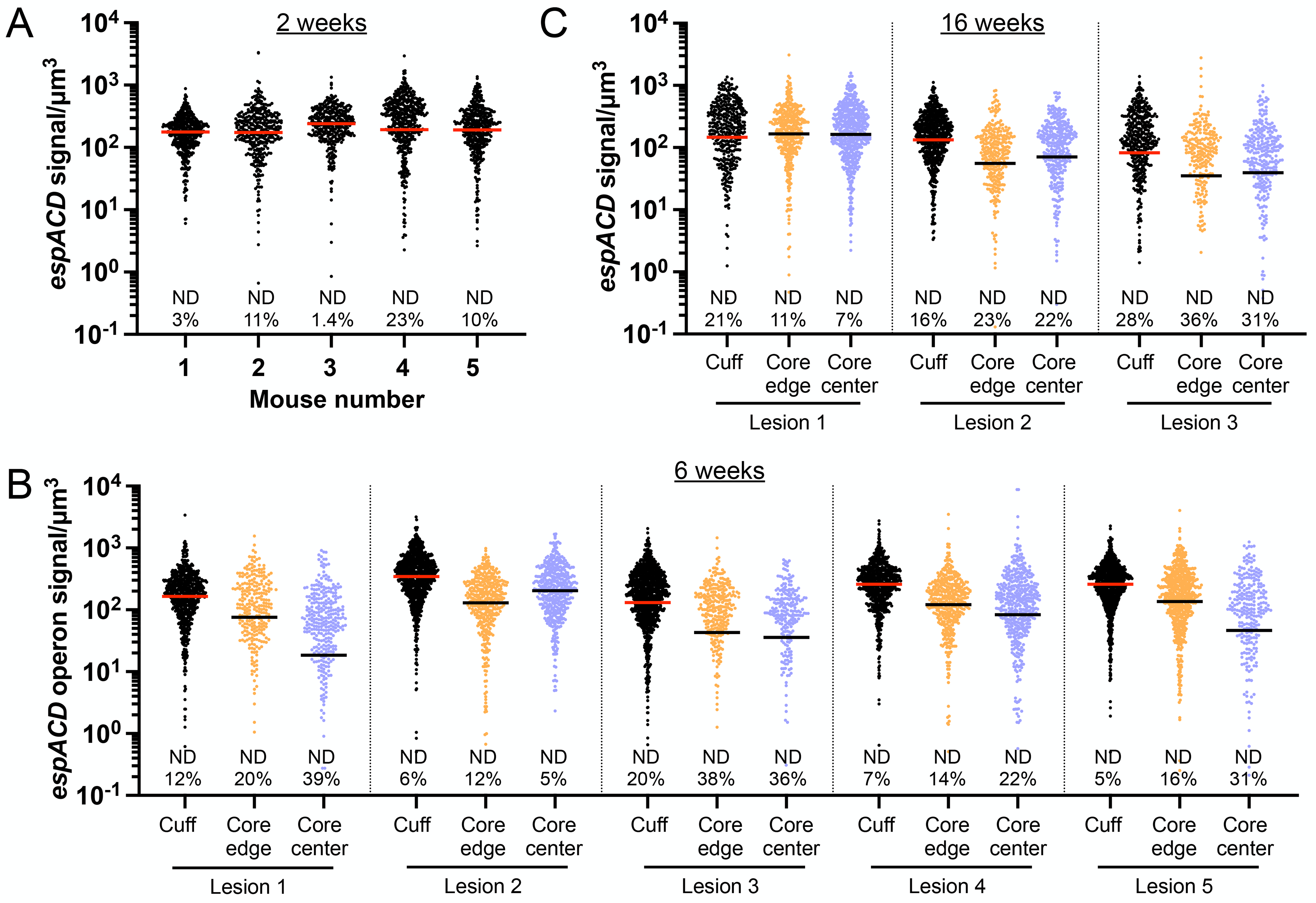
